# Assessment of fish spermatogenesis through high-quality immunofluorescence against fish testicular samples with an antibody set

**DOI:** 10.1101/2022.08.23.504910

**Authors:** Ding Ye, Tao Liu, Yongming Li, Yonghua Sun

## Abstract

A complete evaluation of the spermatogenetic status of a fish by accurately identifying different types of spermatogenic cells is useful not only for reproductive studies but also for genetic breeding. For this task, it is required to establish a simple and practical experimental procedure, to obtain repeatable, high-quality imaging data. Here, we have developed antibodies against the zebrafish (*Danio rerio*) spermatogenesis-related proteins, including Ddx4, Piwil1, Sycp3, and Pcna, and an integrated method for high-quality and high-through output immunofluorescence on testis sections of different fish species. We accurately identified different spermatogenic cells at different stages. Type-A spermatogonia can be identified by the highest expression of Ddx4 and Piwil1 among different spermatogenic cells. Type-B spermatogonia is identified by the second highest expression of Ddx4 and the highest expression of Pcna among different spermatogenic cells. Spermatids can be distinguished from spermatozoa by the expression of Piwil1. The different subtypes of primary spermatocytes (SPC-I) can be identified by co-staining of Sycp3 and Pcna. Leptotene SPC-I show polar expression of both Sycp3 and Pcna at the same side of the nucleus. All the antibodies were tested for practicality in four fish species, Chinese rare minnow (*Gobiocypris rarus*), common carp (*Cyprinus carpio*), blunt snout bream (*Megalobrama amblycephala*), and rice field eel (*Monopterus albus*). Using this method and the antibody sets, we were able to precisely and accurately evaluate the spermatogenetic status in different fish species.

**Highlights:**

1. A practical method for high-quality immunofluorescence against fish testicular samples has been developed.
2. An antibody set of Ddx4, Piwil1, Sycp3 and Pcna have been developed for identification of a variety of spermatogenic cells in different fish species.
3. Antibodies against zebrafish proteins have been tested in the four fish species, Chinese rare minnow, common carp, blunt snout bream and rice field eel.

## 1. Introduction

In various fish species, both genetic and environmental factors, such as sex determine genes and larval nutrition have significant impacts on gametogenesis and reproduction (Helene and Sydney, 2018). Spermatogenesis is a complex process of cellular transformation which produces different types of spermatogenic cells ranging from spermatogonial stem cells (SSCs) to matured sperm (Wang et al., 2022). During fish spermatogenesis, spermatogonia can be further classified into several subtypes, such as type A spermatogonia (SPG-A), type B spermatogonia (SPG-B), while the spermatocytes can be classified into primary spermatocytes of preleptotene (SPC-Pl); leptotene/zygotene (SPC-L/Z); pachytene (SPC-P); diplotene (SPC-D), and secondary spermatocytes (SPG-II) (Schulz et al., 2010; Ye et al., 2019). In the application of surrogate reproduction by spermatogonial transplantation, only undifferentiated SPG-A, also known as spermatogonial stem cells, could be efficiently differentiate into mature gametes in the host fish (Iwasaki-Takahashi et al., 2020; Wang et al., 2022; Xie et al., 2020). Therefore, evaluation of spermatogenetic status of a fish by accurate identifying and quantifying different types of spermatogenic cells, is a very important work not only for fish reproductive research (Ye et al., 2022; Zhang, Q. et al., 2020), but also for germ cell-based genetic breeding such as surrogate reproduction (Jin et al., 2021; Zhang et al., 2022; Zhang, F. et al., 2020).

Characterization of different types of spermatogenic cells is the prerequisites for their quantification, which serves as an accurate method of the phenotypic evaluation of spermatogenetic status. Nuclear characteristic is the most used indicator to characterize different spermatogenic cells (Leal et al., 2009). However, this method requires high quality images of nuclear staining of the entire section of testis samples, and thus a reliable histology method to preserve nuclear morphology is needed. Moreover, nuclear-based identification also highly relies on research experience of nuclear morphology of specific cell type.

To compensate the difficulties of nuclear characterization, many molecular markers were identified and some of their antibodies were developed to characterize the spermatogenic cells especially the spermatogonial stem cells in fish (Lacerda et al., 2014). Ddx4 (previously named Vasa), a DEAD-box family member with RNA helicase activity, plays critical roles in germ granule formation and PIWI-interacting RNAs (piRNAs) biogenesis (Thomson and Lin, 2009; Xiol et al., 2014; Xu et al., 2021). Piwil1 (preciously name Ziwi), which belongs to the Argonaute protein family, involves piRNA biogenesis and silencing of transposable elements (Ponnusamy et al., 2017). Both Ddx4 and Piwil1 are used as the classical germ cell markers. The germline expression pattern of Ddx4 has been reported in human (Castrillon et al., 2000), and several species in mammals (Kim et al., 2015; Toyooka et al., 2000), reptile (Liu et al., 2021) and fish (Cao et al., 2012; Leu and Draper, 2010; Nobrega et al., 2010; Xu et al., 2005). The germline expression pattern of Piwil1 has been reported in *Drosophila* (Gonzalez et al., 2015), zebrafish (Houwing et al., 2008; Leu and Draper, 2010; Ye et al., 2019), Nile tilapia (Xiao et al., 2013), mice (Deng and Lin, 2002), and human (Qiao et al., 2002). Sycp3 (alternative name SCP3) is one of the lateral element proteins of the synaptonemal complex which forms during homologous chromosome synapsis (Biswas et al., 2021). It has been used as a marker for spermatocyte at different stages of meiotic prophase in zebrafish (Ozaki et al., 2011), medaka (Iwai et al., 2006), and mice (Parra et al., 2004). Proliferating cell nuclear antigen (Pcna) is a component of the DNA replication and repair machinery helping DNA polymerases binding to the DNA (Gonzalez-Magana and Blanco, 2020). It has been widely used as a marker for cell proliferation (Cardano et al., 2020; Jurikova et al., 2016). Antibodies against the above proteins would be useful as the markers to precisely evaluate the spermatogenetic status in fish. However, there lacks a set of reliable antibodies against fish derived proteins, and it is also unknown whether the antibody against a certain fish species could be used in other fish species.

In this study, we not only generated a set of antibodies against zebrafish spermatogenesis-related proteins, but also developed an integrated method for high quality immunofluorescence on testicular sections of different fish species by using these antibodies.

## 2. Materials and methods

### 2.1. Fish

One-year old zebrafish (*Danio rerio*) of the AB strain and one-year old wild-type rare minnow (*Gobiocypris rarus*) were obtained from the China Zebrafish Resource Center, National Aquatic Biological Resource Center (CZRC-NABRC, http://zfish.cn). Two-year old common carp (*Cyprinus carpio*) and two-year old blunt snout bream (*Megalobrama amblycephala*) were obtained from the NABRC, Institute of hydrobiology, Chinese Academy of Sciences as previously described (Zhu et al., 2019). Rice field eel (*Monopterus albus*) was bought from a local fish market.

### 2.2. Generation of polyclonal antibodies

A custom antibody production service from the company (Bioyear, China) was utilized to generate rabbit polyclonal antibodies. The epitope of Piwil1 was EGQLVGRGRQKPAPGC according to a previous study (Houwing et al., 2007). The epitopes of Ddx4, Sycp3 and Pcna were designed to cover the conserved region of certain protein respectively, and the epitope sequences were listed in the Table 1. Briefly, zebrafish protein sequences were applied to protein blast analysis on NCBI (https://blast.ncbi.nlm.nih.gov). Based on the top 100 hits, the average identity between zebrafish and other fish was above 65% for Sycp3, 80% for Piwil1, 85% for Ddx4, and 95% for Pcna, and the conserved regions were identified. All the antibodies were purified from anti-serum by using antigen-affinity chromatography. The antibodies have been deposited to CZRC-NABRC for distribution within research community.

### 2.3. Antibody labeling with fluorescence

For the co-staining of Pcna and Sycp3, the antibodies were labeled with Alexa Fluor® 568 and Alexa Fluor® 488 respectively using the Alexa Fluor® Antibody Labeling Kits according to the manufacturer’s manual (Thermo Fisher). The antibodies for fluorescence labeling were dissolved in PBS without any glycerol.

### 2.4. Image-based evaluation of spermatogenesis

A standard image-based evaluation of spermatogenesis comprises the following steps: (1) preparation of adhesive cover glass, (2) sample preparation and frozen section, (3) immunofluorescence, (4) sample mounting and confocal imaging, (5) image processing and analysis. A simplified work flow is illustrated in Figure 1.

**Figure 1.**
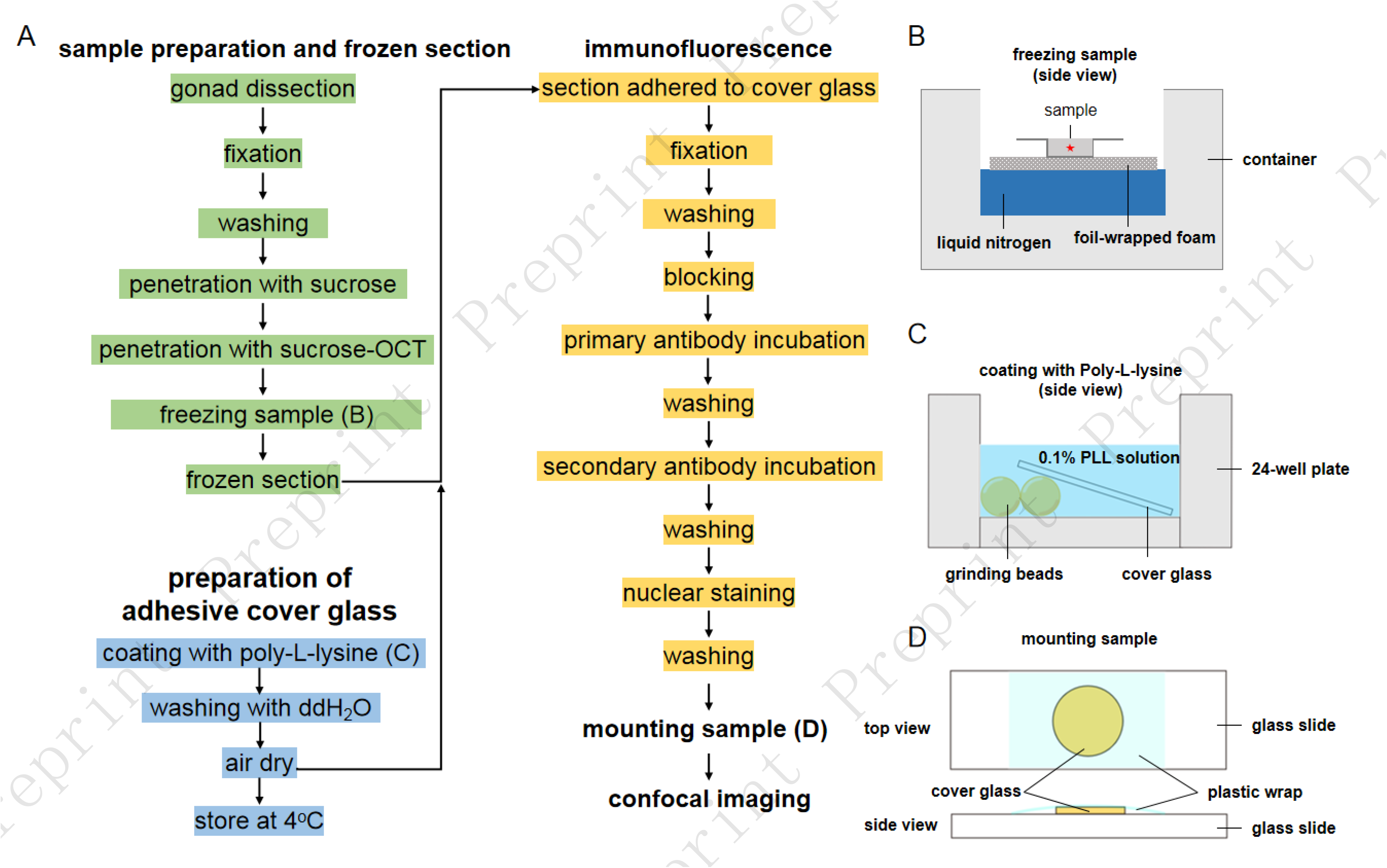
A simplified work flow of image-based evaluation of spermatogenesis. (A) Route diagram of the experimental procedures; (B) Schematic showing freezing sample in OCT; (C) Schematic showing coating with PLL; (D) Schematic showing mounting sample.

#### 2.4.1. Preparation of adhesive cover glass

In order to achieve high-throughput immunofluorescence, 24-well plates were used for the following manipulations. To facilitate tissue sections adherence, round cover glasses with diameter of 12 mm were coated with Poly-L-lysine (PLL) and evaluated as previously described (Liu et al., 2022). Briefly, eight zirconia grinding beads were placed in each well, a cover glass was put on them, and 500 μL PLL solution (1 mg/mL, Sigma-Aldrich) was added to immerse the cover glass (Figure 1 B). After 30 min, the PLL solution was removed and stored at 4°C for reuse. One mL of ddH_2_O was added to each well to wash the cover glass for 5 min, and the PLL-coated cover glasses were air-dried in an oven at 60 °C. The 24-well plate containing adhesive cover glasses was sealed with plastic wrap and stored at 4°C for use.

#### 2.4.2. Sample preparation and frozen section

Fish were anesthetized with 0.2 mg/mL MS-222 in fish system water and the testes were dissected out. For zebrafish, the testis was fixed in 4% PFA at room temperature (RT) (~25 °C) for 2 hours. The fixed samples were processed for frozen section according to a previously published method with modifications (Barthel and Raymond, 1990). Briefly, the fixed testis were wash with pH 7.0 PBST buffer (PBS with 0.1% trionX-100) for 3 times of 5 min, and transferred into a series of sucrose in PBS solution for 30 min each: 5% sucrose; 8% sucrose; 12% sucrose;16% sucrose; 20% sucrose until tissue sinking to the bottom. Samples were washed in 20% sucrose/OCT (2:1) and 20% sucrose/OCT (1:1) each for 30 min, and finally embedded in 20% sucrose/OCT (1:1) in a mold on foil-wrapped floating foam on liquid nitrogen (Figure 1 C). The embedded testis was cut into 12 μm, and the sections were collected to a PLL-coated cover glasses.

#### 2.4.3. Immunofluorescence

The sections were air dried for 15 min, and fixed with 4% PFA at RT for 20 min. After washing with PBST (0.1% tritonX-100 in PBS) for 3 times of 5 min, the sections were permeabilized by 0.5% Triton X-100 in PBS for 30 min, and washed with PBSBDT (2% BSA, 1% DMSO, 0.1% tritonX-100 in PBS) for 3 times of 5 min. The sections were then blocked in PBSBDT for 1 hour at RT. After blocking, the sections were incubated in the diluted antibodies (1:200 in PBSBDT) overnight at 4 °C. After washing with PBSBDT for 3 times of 5 min, the sections were incubated in secondary antibody (Goat-anti-Rabbit Alexa Fluor 488, 1 mg/mL in stock, 1:500 in PBSBDT) (Thermo Fisher) at RT for 5 hours. The sections were then washed with PBSBDT for 3 times of 5 min to remove the unbound antibody, stained with DAPI at 1μg/mL at RT for 30 min, and washed with PBST for 3 times of 5 min again. For the co-stain of Pcan and Sycp3, the incubation of secondary antibody and the following wash steps were omitted.

#### 2.4.4. Sample mounting and confocal imaging

After immunofluorescence, each section was mounted with 10 μL VECTASHIELD® mounting medium, and covered with a piece of plastic wrap (Figure 1 D). The samples were stored at 4°C in a dark box till applying for confocal microscopy. Confocal images were acquired using a laser-scanning confocal inverted microscope (SP8, Leica) with a 20× air objective or a 63× oil-immersed objective.

#### 2.4.5. Image processing and analysis

Images were adjusted with brightness and contrast with Fiji software (Schindelin et al., 2012).

## 3. Results

### 3.1. Identification of SPG-A, SPG-B, SPC-I, SPD and SPZ based on expression of Ddx4

By using the developed method, we could obtain high-quality immunofluorescence results with fish testicular samples (Figure 2–10). In zebrafish testis, SPG-A, SPG-B, SPC-I, SPD and SPZ can be identified according to the expression pattern of Ddx4 (Figure 2 A-A1). Ddx4 was found with highest expression level in SPG-A (Figure 2 arrow in A1; B-B2; C-C2), and second highest in SPG-B (Figure 2 D-D2). As SPG-B differentiating to SPC-I, the expression level of Ddx4 was dramatically reduced (Fig 2 E-E2). The SPD showed very weak expression of Ddx4, while the spermatozoa show no expression (Figure 2 F-F2).

**Figure 2.**
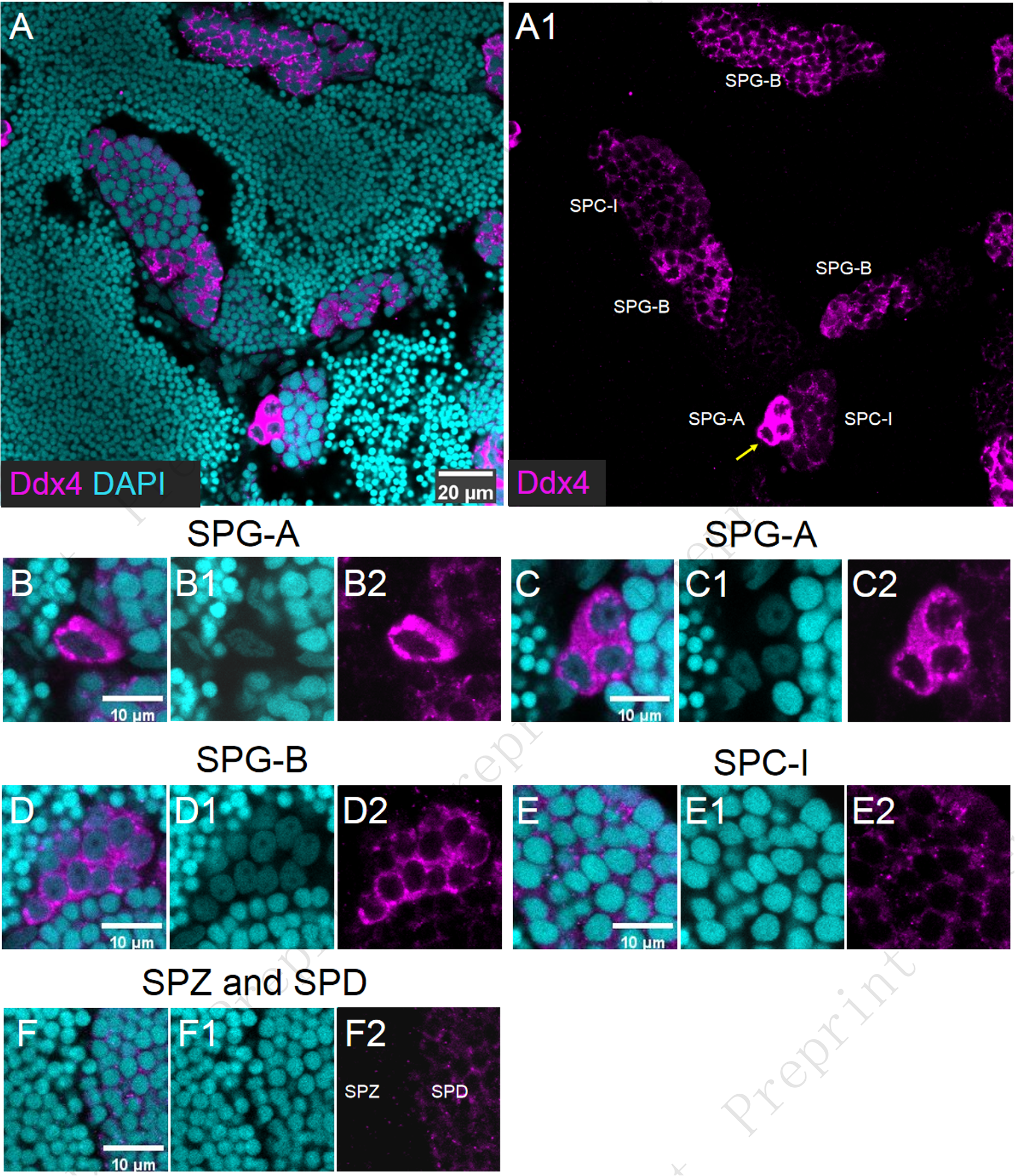
Identify SPG-A, SPG-B, SPC-I, SPD and SPZ based on expression of Ddx4. (A-A1) Representative images of a zebrafish testis section showed the expression of Ddx4 (Magenta) co-stained with DAPI (Cyan). The arrow pointed out SPG-A (A1). Representative images showed single-located SPG-A (B-B2), cyst SPG-A (C-C2), SPG-B (D-D2), SPC-I (E-E2), SPD and SPZ (F-F2). The single-located SPG-A and cyst SPG-A was detected with the highest level of Ddx4. The SPG-B was detected with the second highest level of Ddx4. The expression level of Ddx4 was dramatically reduced in SPC-I compared to that in SPG-B. The SPD showed very weak expression of Ddx4, while the spermatozoa show no expression.

In addition, the nuclear characteristics of SPG-A, SPG-B and SPC-I can be used as secondary indicator. Although both SPG-A and SPG-B have weak DNA staining, SPG-A is usually found with one large nucleolus, while the SPG-B with one or two smaller nucleolus (Figure 2 C1, D1). SPC-I has higher DNA staining intensity than SPG, which make it distinguishable (Figure 2 E1). Taken together, simultaneously considering the nuclear characteristics and Ddx4 expression pattern, we could identify SPG-A, SPG-B, SPC-I, SPD and SPZ in zebrafish.

The antibody against zebrafish Ddx4 also worked perfectly in testicular samples of Chinese rare minnow, common carp, blunt snout bream, and rice-field eel. All these fish showed similar expression pattern of Ddx4 as zebrafish (Figure 3 A-D; A1-D1). Notably, in the testis of rice field eel, distinguishable expression of Ddx4 could be seen in the SPG-A, but not in SPG-B or SPC-I, suggesting Ddx4 could serve as a marker for SPG-A of rice field eel (Figure 3 D and D1).

**Figure 3.**
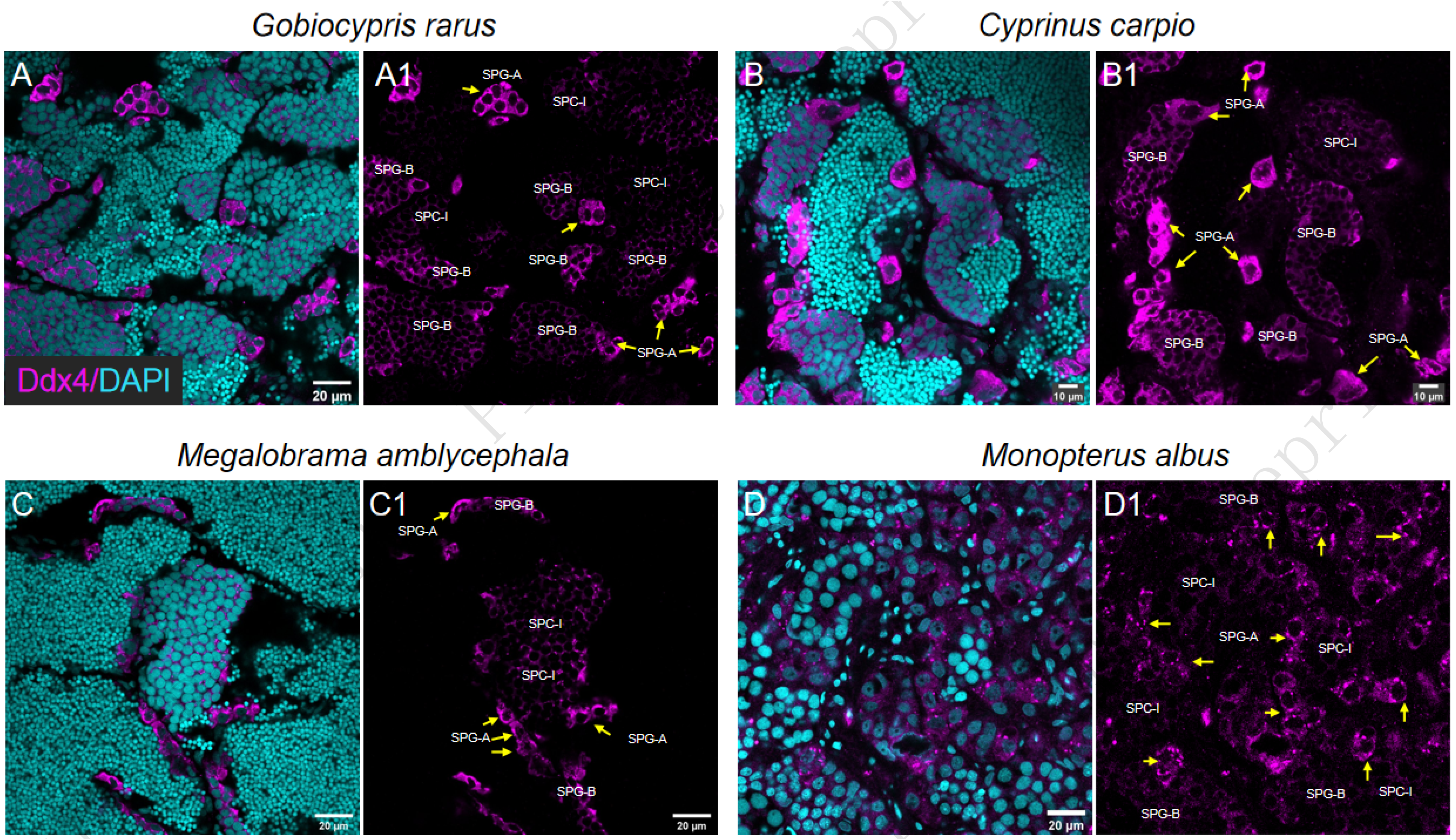
Ddx4 expression pattern in other fish species. Representative images show the expression of Ddx4 in *Gobiocypris rarus* (A, A1), *Cyprinus carpio* (B, B1), *Megalobrama amblycephala* (C, C1), *Monopterus albus* (D, D1). The arrows pointed out SPG-A.

### 3.2. Identification of SPG-A, SPD and SPZ based on expression of Piwil1

In zebrafish testis, Piwil1 was found with highest expression level in SPG-A (Figure 4 A-A1; B-B2). A comparable expression of Piwil1 was found in SPG-B and SPC-I (Figure 4 A-A1), which makes it impossible to identify these two cell types solely based on Piwil1 expression. The characteristics of nuclear staining of SPD and SPZ were indistinguishable (Figure 4 C1), and Piwil1 was remarkably expressed in SPD but not in SPZ, suggesting Piwil1 is a good Indicator to identify the two cell types (Figure 4 A1; C-C2). Thus, Piwil1 can be used to identify SPG-A, SPD and SPZ in zebrafish.

**Figure 4.**
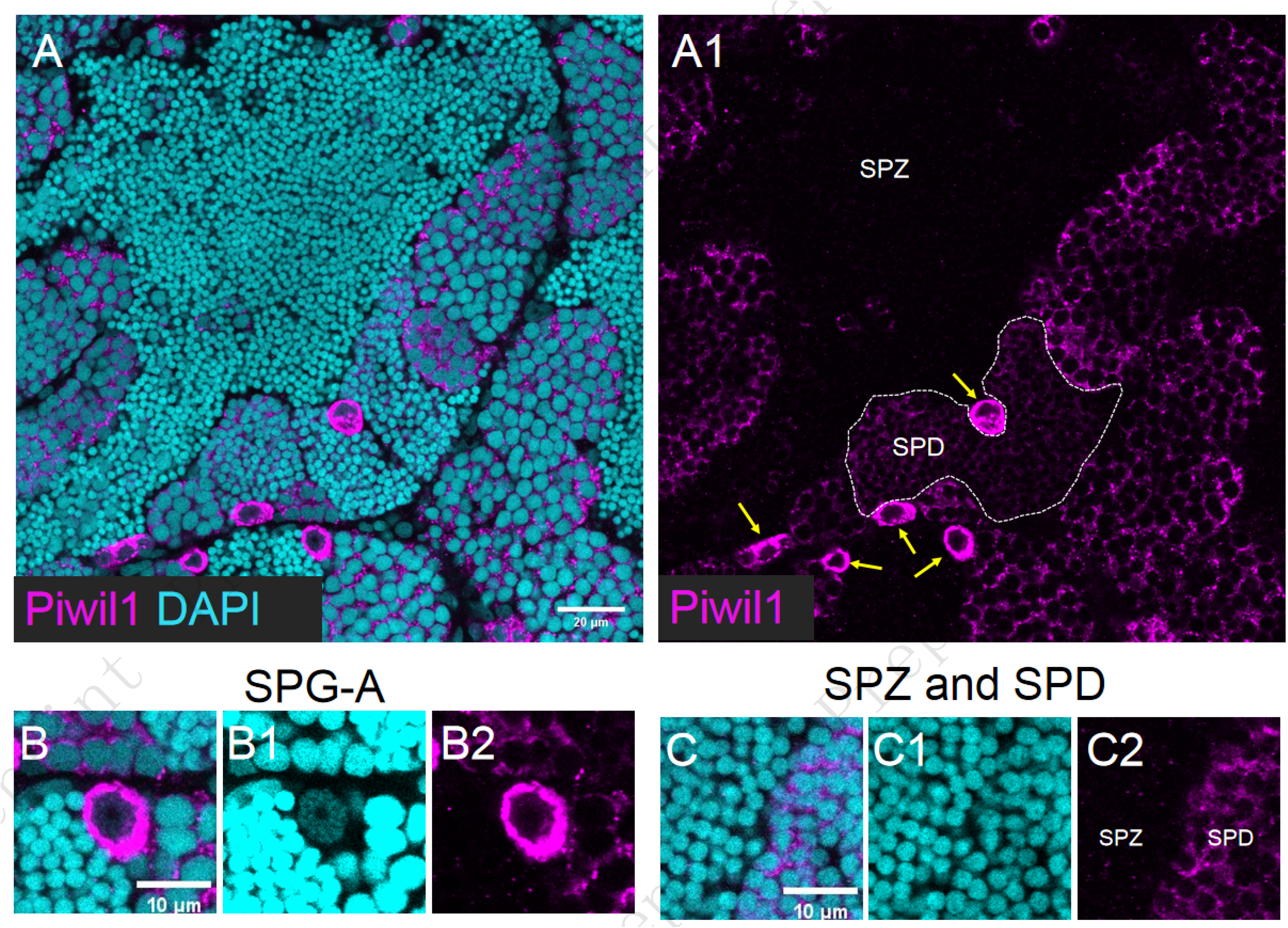
Identify SPG-A, SPD and SPZ based on expression of Piwil1. (A-A1) Representative images of a zebrafish testis section showed the expression of Piwil1 (Magenta) co-stained with DAPI (Cyan). The arrows pointed out SPG-A (A1). Representative images showed the single-located SPG-A (B-B2), SPD and SPZ (C-C2). Piwil1 was found with highest expression level in SPG-A.

The antibody against zebrafish Piwil1 also worked in testicular samples of Chinese rare minnow, common carp, and blunt snout bream, but not of rice field eel (Figure 5 and not shown). However, unlike in zebrafish, the expression level of Piwil1 was undistinguishable between SPG-A and SPG-B in these three fish (Figure 5).

**Figure 5.**
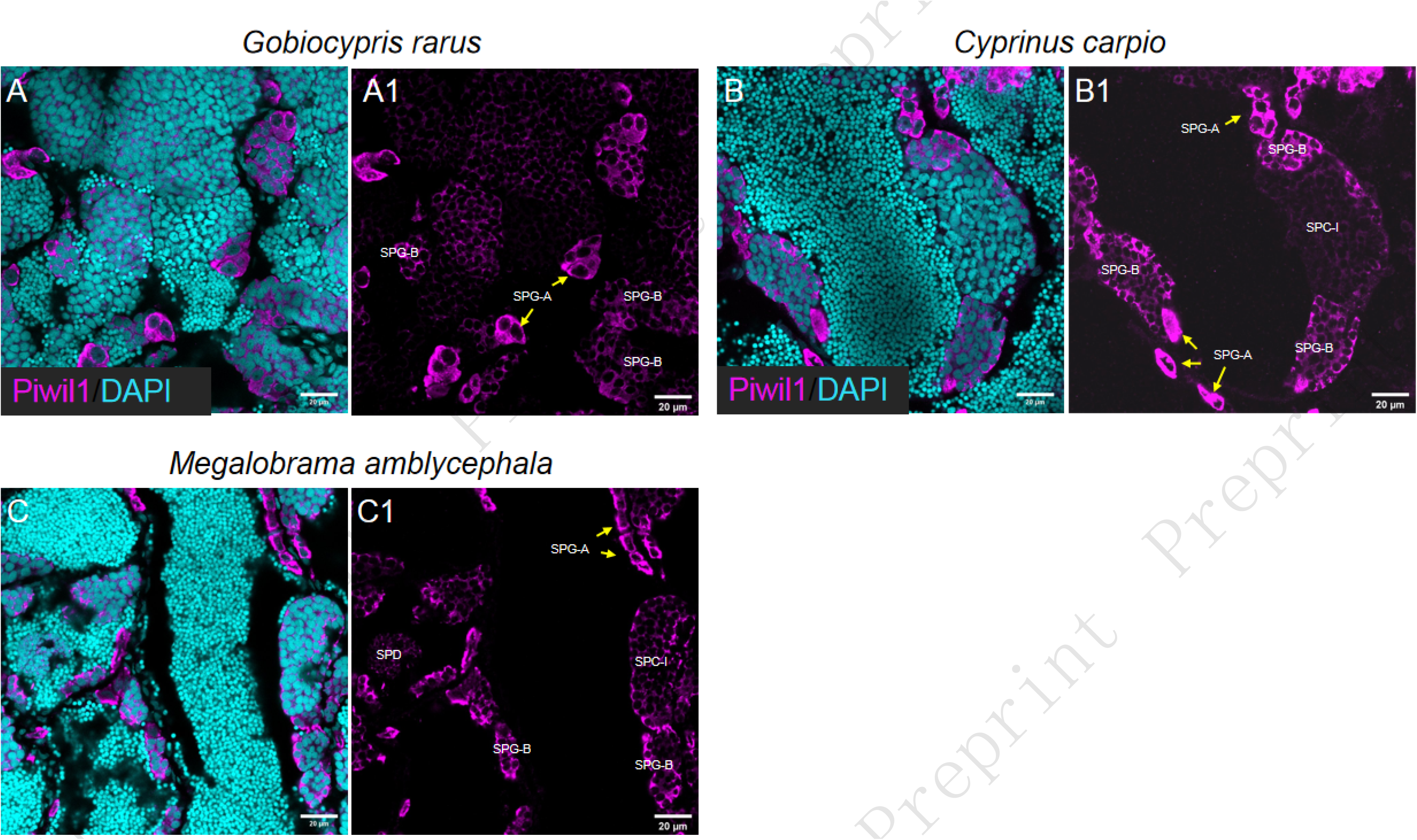
Piwil1 expression pattern in other fish species. Representative images showed the expression of Piwil1 in *Gobiocypris rarus* (A, A1), *Cyprinus carpio* (B, B1), *Megalobrama amblycephala* (C, C1). The arrows pointed out SPG-A.

### 3.3. Identification of SPG and SPC-II based on expression of Pcna

In zebrafish testis, Pcna was highly expressed in both SPG-A and SPG-B, but lowly expressed in SPC and SPD, suggesting that Pcna can be used as marker for SPG (Figure 6 A-A1; B-B2; C-C2). Interestingly, Pcna was found polarly localized in the nucleus of a subtype of SPC-I (Figure 6 D-D2) which was identified as SPC-I at leptotene stage in the following study, and it was also found highly expressed in SPC-II (Figure 6 E-E2). Thus, Pcna can be used to identify SPG-A and SPC-II in zebrafish.

**Figure 6.**
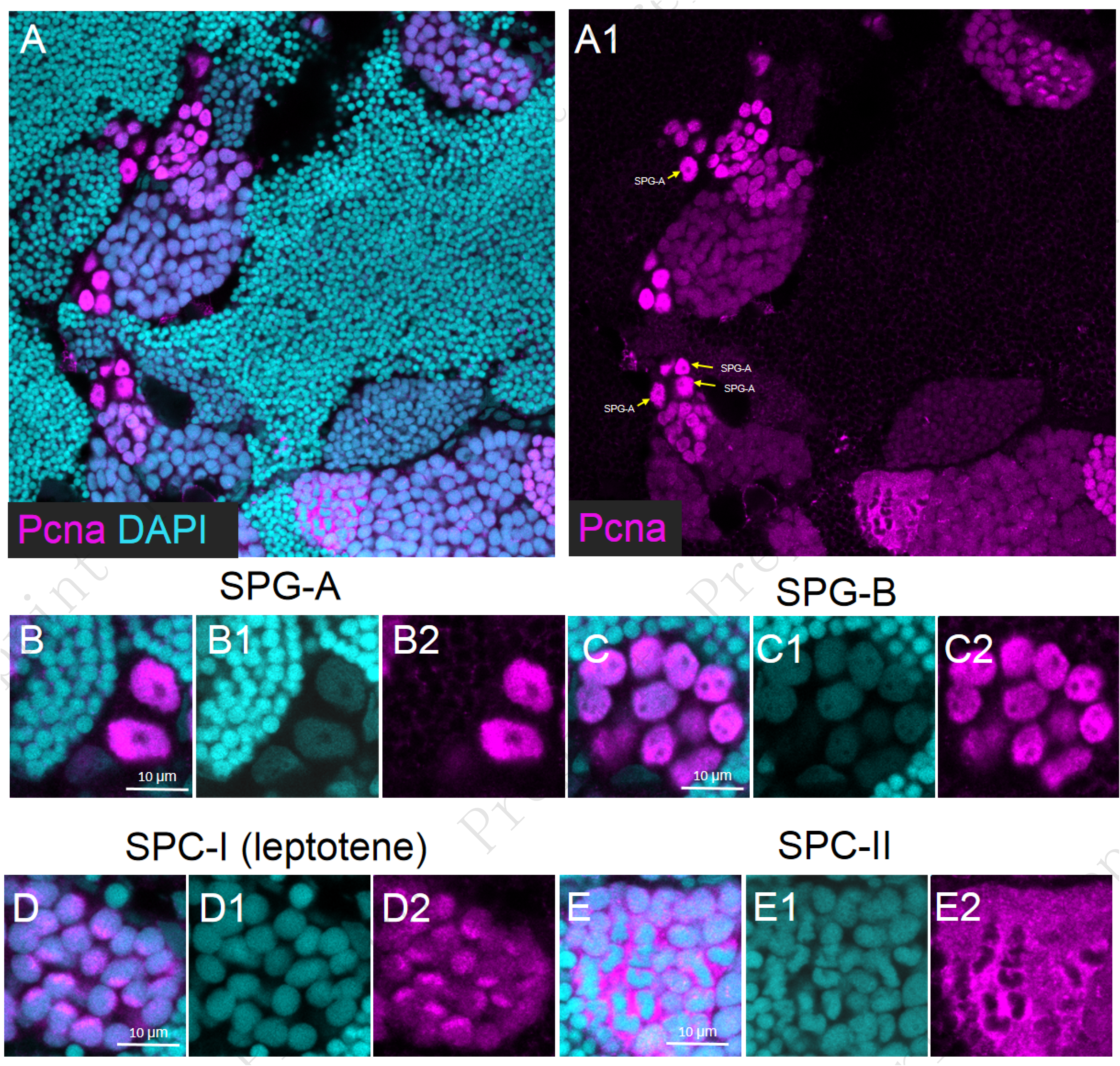
Identify SPG and SPC-II based on expression of Pcna. (A-A1) Representative images of a zebrafish testis section showed the expression of Pcna (Magenta) co-stained with DAPI (Cyan). The arrow pointed out SPG-A (A1). Representative images showed the SPG-A (B-B2), SPG-B (C-C2), SPC-I at leptotene stage (D-D2), SPC-II (E-E2). Pcna was highly expressed in both SPG-A, SPG-B and SPC-II and polarly localized in the nucleus SPC-I at leptotene stage.

The antibody against zebrafish Pcna also worked in the four different fish as mentioned above (Figure 7). The expression pattern of Pcna in testis is similar between zebrafish and Chinese rare minnow (Figure 7 A and A1). We also found ploarly nuclear localized-Pcna in the SPC-I of the Chinese rare minnow (Figure 7 A1). Interestingly, unlike zebrafish, the expression of Pcna was restricted in the cytoplasm of the SCP-I in the common carp (Figure 7 B and B1). Moreover, we did not find ploarly nuclear localized-Pcna in the SPC-I of blunt snout bream and rice field eel, and this might be due to the developmental status of the testis in those fish.

**Figure 7.**
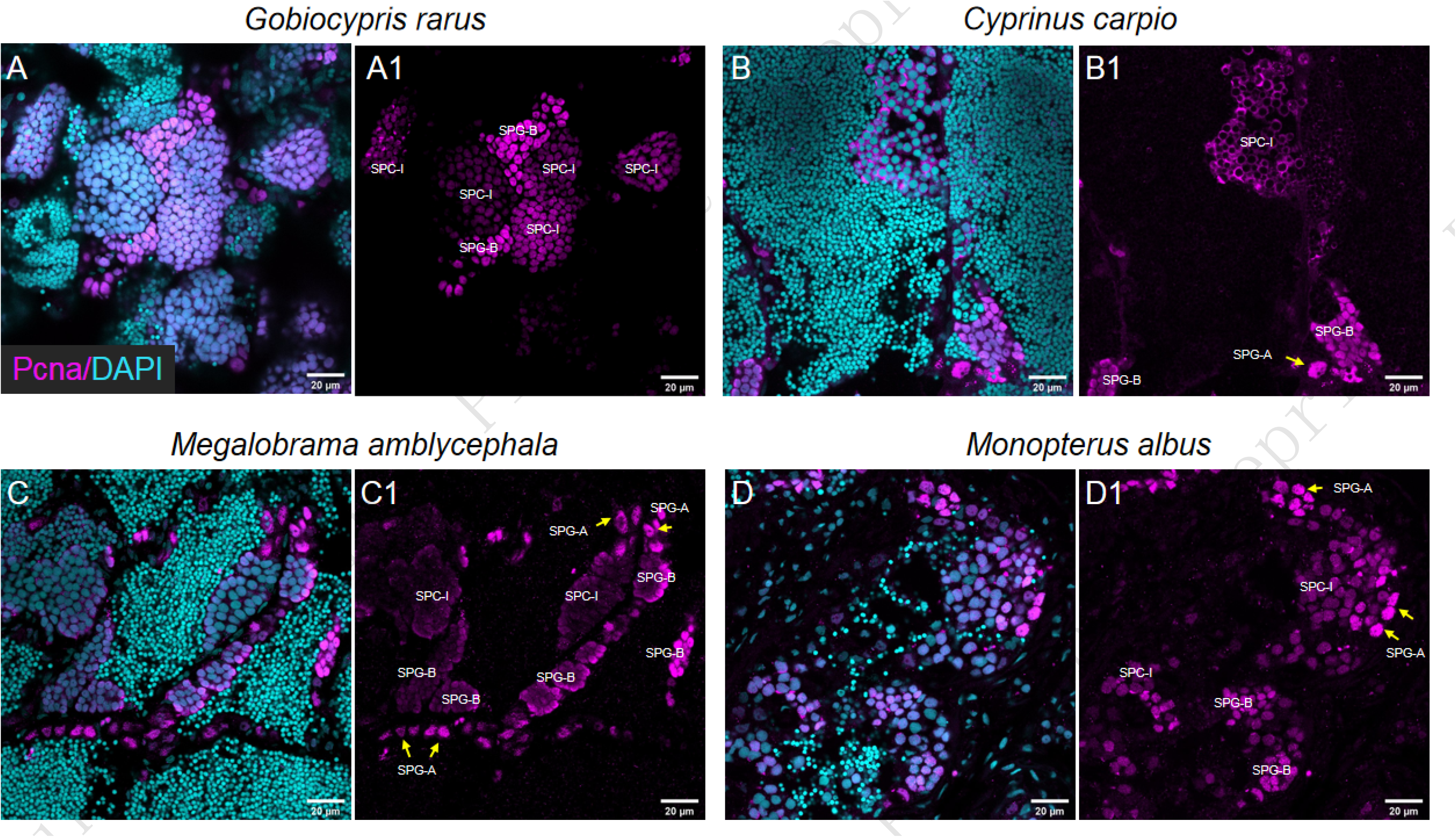
Pcna expression pattern in other fish species. Representative images showed the expression of Pcna in *Gobiocypris rarus* (A, A1), *Cyprinus carpio* (B, B1), *Megalobrama amblycephala* (C, C1), *Monopterus albus* (D, D1). The arrows pointed out SPG-A.

### 3.4. Identification of SPG-A, SPG-B and different subtypes of SPC-I based on co-expression of Sycp3 and Pcna

In this section, the different subtypes of SPC during meiosis I were staged according to the previous article (Imai et al., 2021). Sycp3 was found with unique expression pattern in different subtypes of SPC-I. The expression of Sycp3 was low in the SPG, polarly localized in the nucleus of leptotene SPC-I, distributed within the whole part of nucleus at pachytene stage, and dramatically reduced in the nucleus of diplotene SPC-I (Figure 8 A-A1; B-B2; C-C2; D-D2). We also tested this antibody in the four fish and found it only worked in rice field eel (Figure 9).

**Figure 8.**
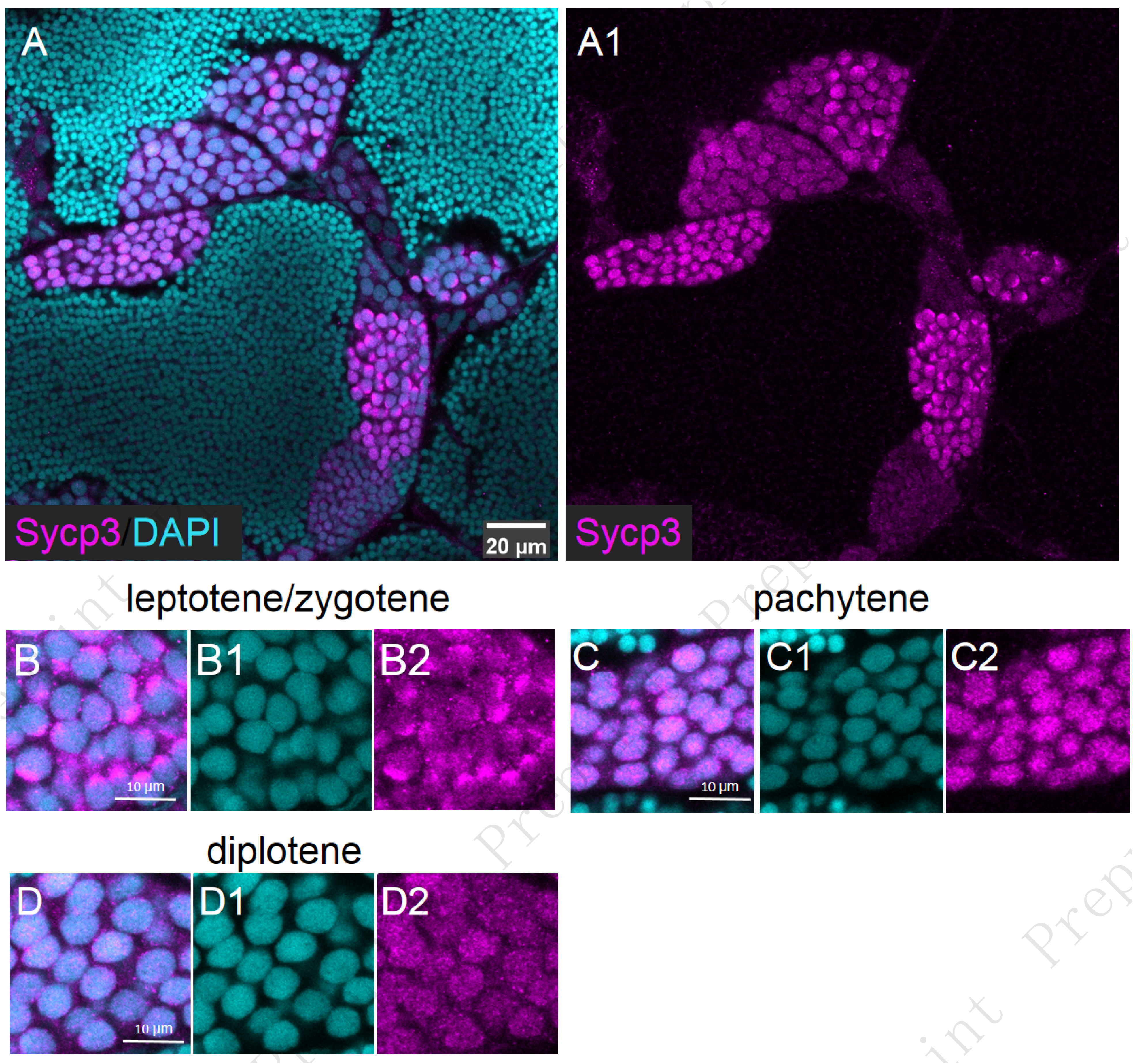
Identify different subtypes of SPC-I based on expression of Sycp3. (A-A1) Representative images of a zebrafish testis section showed the expression of Sycp3 (Magenta) co-stained with DAPI (Cyan). (B-B2) Representative images of SPC-I at leptotene stage; (C-C2) Representative images of SPC-I at pachytene stage; (D-D2) Representative images of SPC-I at diplotene stage.

**Figure 9.**
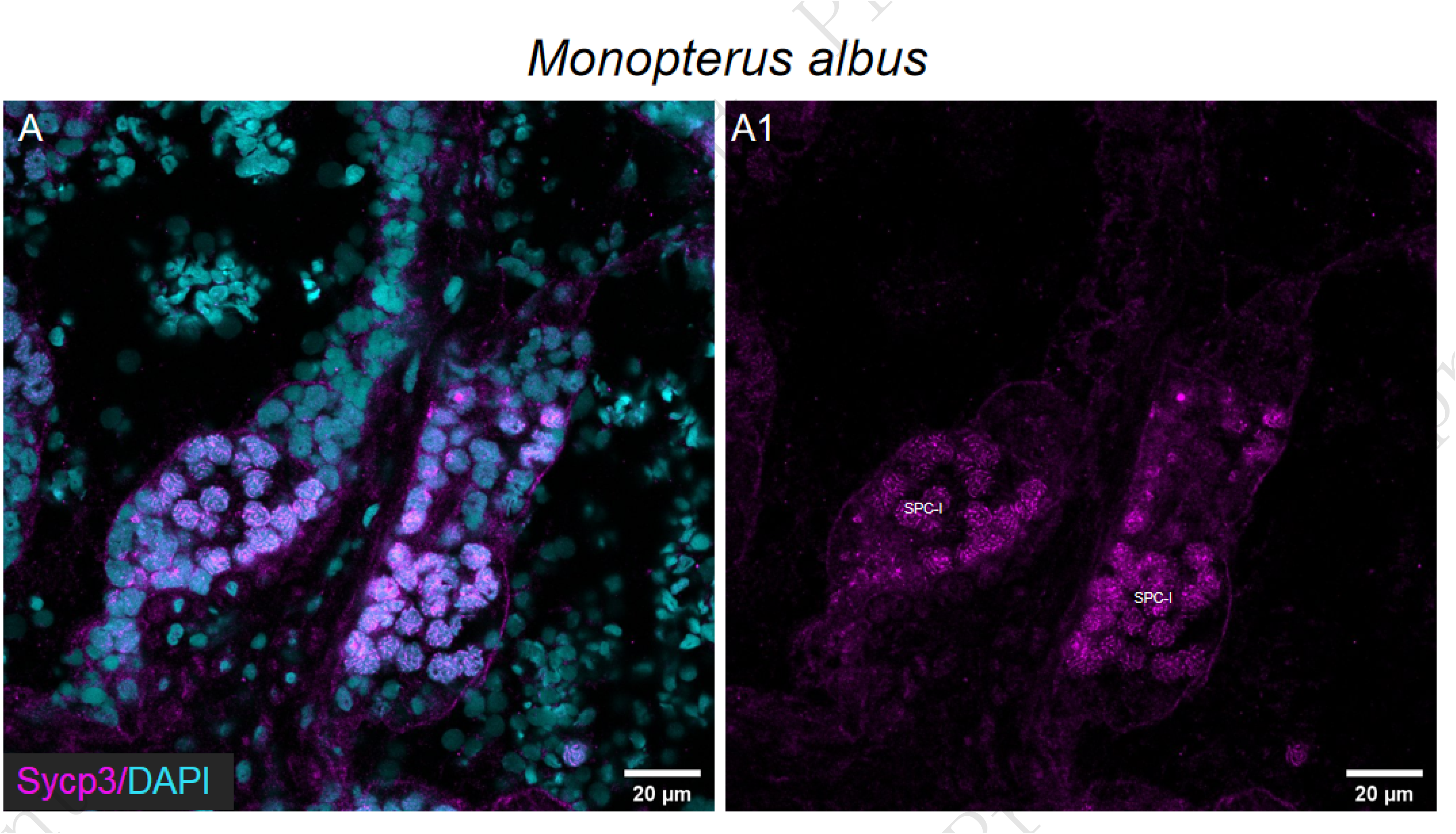
Sycp3 expression pattern in other fish species. Representative images showed the expression of Sycp3 in *Monopterus albus* (A, A1).

Since both Pcna and Sycp3 have unique expression patterns in the spermatogenic cells, we co-stained Pcna and Sycp3 and applied to lightening confocal imaging to monitor the developmental changes of spermatogenesis from SPG to SPC-I. Both SPG-A and SPG-B had Pcna expressed but not Sycp3 (Figure 10 A, B). Notably, the expression the Pcna was not identical among all the SPG-A cells, which raise the possibility that Pcna expression level of a cell is positive-correlated with its mitosis ability (Figure 10 A). When the Sycp3 showed polarly localized in the nucleus and Pcna expression decreased, the cells entered from SPG-B stage into SPC-preleptotene stage (Figure 10 C). At leptotene stage, both Sycp3 and Pcna polarly localized at the same side in the nucleus with Sycp3 more marginally located (Figure 10 D). As SPC-I entering from leptotene to zygotene stage, the expression of polarly located Sycp3 increased while Pcna lost its polarity and the expression level was decreased (Figure 10 E). At mid - pachytene stage, the expressional region of polarly located Sycp3 extended to nearly half of the nucleus with parallel strip pattern while Pcna remained at low level (Figure 10 F). At late - zygotene stage, Sycp3 expressed at the whole part of the nucleus with parallel strip pattern while Pcna displayed punctated expression pattern (Figure 10 G). At pachytene stage, the strip pattern of Sycp3 changed from parallel to interlace and the expression level decreased, while Pcna was nearly undetectable (Figure 10 H). It is difficult to identify SPC at diplotene stage. Therefore, the different subtypes of SPC-I as well as SPG could be distinguished through co-staining of Sycp3 and Pcna.

**Figure 10.**
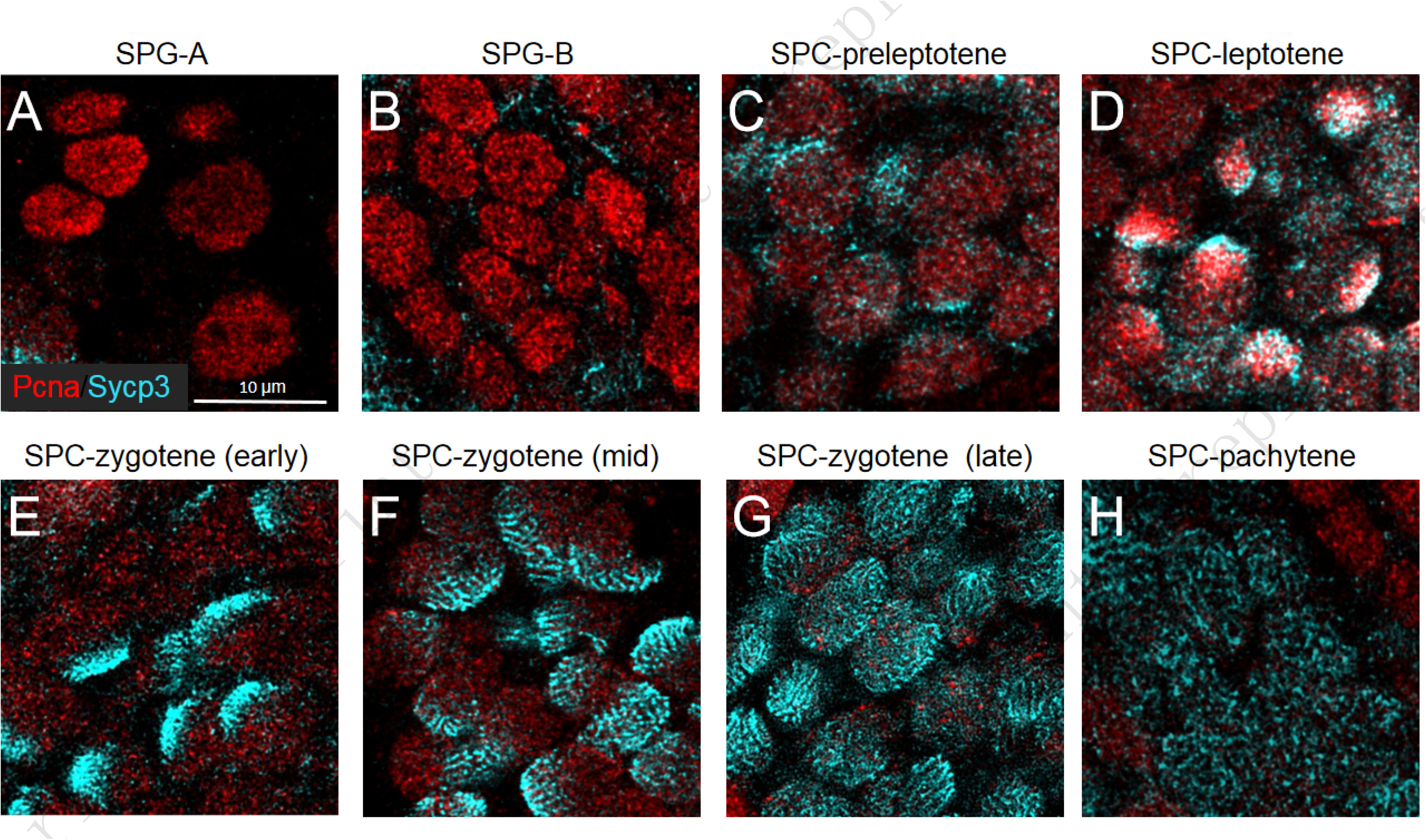
Identify SPG-A, SPG-B and different subtypes of SPC-I based on co-expression of Sycp3 and Pcna. Representative image showing SPG-A (A), SPG-B (B), preleptotene SPC (C), early leptotene SPC (D), late leptotene SPC (E), early pachytene SPC (F), late pachytene SPC (G), diplotene SPC (H).

## 4. Discussion

In this study, we have improved immunofluorescence in the following aspects. 1) To archive high throughput by using 24-well plate, which makes it possible to perform the experiment on an automatic pipetting station; 2) The coating protocol for the preparation of adhesive cover glasses is provided. Although, there are commercial adhesive cover glasses available on the market, home-made ones always give us the best practice. Moreover, the PLL-coated cover glass should be completely dry out and stored avoiding from air and humid; 3) To use 50%OCT-10%sucrose instead of 100% OCT as the embedding media for freezing, as the former one can better maintain the cell morphology than the latter; 4) To freeze sample on a foil-wrapped foam on liquid nitrogen is simple, avoiding the usage of isopentane, a volatile chemical (Douglas and Kenneth, 1990).

In this study, we identified all types of spermatogenic cells in zebrafish using a set of home-made antibodies, which are against Ddx4, Piwil1, Pcna and Sycp3 (Figure 11). All the antibodies have been tested for the applicability in four fish species, Chinese rare minnow, common carp, blunt snout bream, and rice field eel. The Ddx4 and Pcna antibodies work in all the tested fish. The Piwil1 antibody does not work in rice field eel, while the Sycp3 antibody works in rice field eel but not in other three fish species. These data demonstrate that by designing a conserved antigen epitope of certain protein from different fish species, the antibody against zebrafish protein have high possibility working in other fish species.

**Figure 11.**
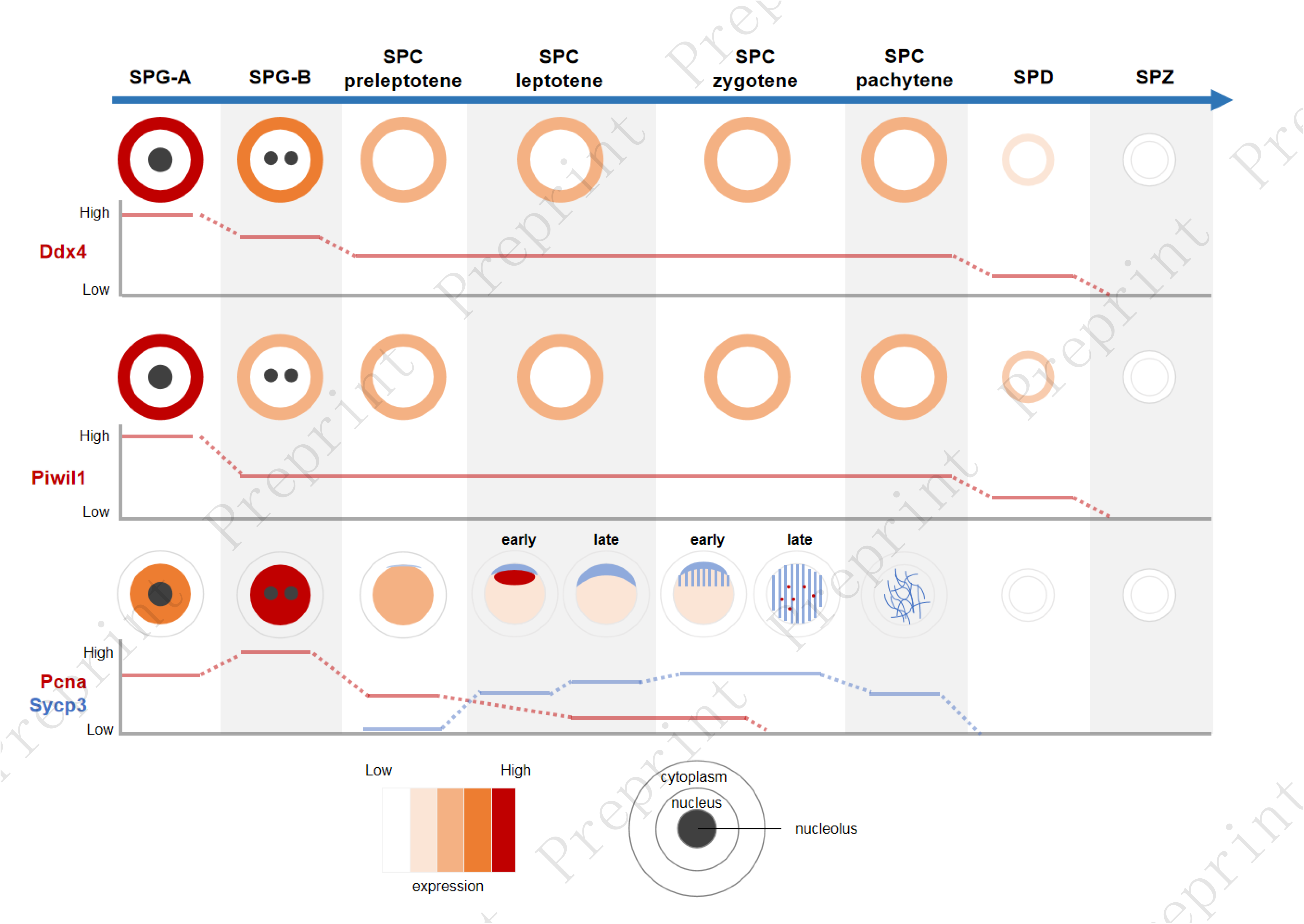
A graphic summary for identification of spermatogenic cell based the expression pattern of Ddx4, Piwil1, Pcna and Sycp3.

The previous and our studies suggest that the expression pattern of Ddx4 in different types of spermatogenic cells is consistent among different fish species, with the highest expression in the spermatogonia, and nearly not detectable in the spermatid (Cao et al., 2012; Kobayashi et al., 2000, 2002; Xu et al., 2005). Interestingly, the *Apostichopus japonicus* also showed similar expression pattern of Ddx4 in the testis to that of fish species (Liu et al., 2021). However, this pattern is different in other vertebrate species. For example, the expression level of Ddx4 was higher in the spermatocyte than that in the spermatogonia and spermatid in human (Castrillon et al., 2000), Asian yellow pond turtle (Liu et al., 2021), and stallions (Kim et al., 2015). This consistent expression signature of Ddx4 expanded the usage of Ddx4 antibody to identify different types of spermatogenic cells for different fish species.

Piwil1 was detected with the highest level in the SPG-A in zebrafish and Chinese rare minnow. However, the pattern is different in common carp and blunt snout bream, in which the expression of Piwil1 between SPG-A and SPG-B are comparable. In all the tested fish species, Piwil1 was detected in SPD but not SPZ. In human, however, the expression of Piwil1 in testis is different, with expression in pachytene SPC-I and SPD but not in SPG (Hempfling et al., 2017). Thus, Piwil1 may not have a common expression pattern in testis among different vertebrates, although Piwil1 is exclusively expressed in germ cells.

In all the tested fish, Pcna has a consistent expression pattern in SPG-A and SPG-B, suggesting it is a reliable marker for the mitotic SPG-A. Interestingly, Pcna were restricted in the cytoplasm of SPC-I in the common carp, whereas it was down-regulated in the SPC-I in other tested fish species, suggesting that Pcna possess different expressional regulation mechanism in different fish. The cytoplasmic-localized Pcna has been found in mature neutrophils with anti-apoptosis function (Bouayad et al., 2012). Other study about acute myeloid leukemia also revealed that the leukemic cells export PCNA to cytoplasm to promote cell survival (Ohayon et al., 2016). Thus, the common carp might have a unique ability to regulate the translocation of Pcna from nucleus to cytoplasm to promote the survival of SPC-I.

It has been reported that Pcna promote meiotic recombination by activating the MutLγ endonuclease (Kulkarni et al., 2020). However, the dynamic expression pattern of Pcna during meiosis is still missing. In this study, we carefully characterized the expression dynamic of Pcna during meiosis I in zebrafish, and found that Pcna was polarly localized in the nucleus of leptotene SPC-I just underneath the Sycp3, suggesting a potential role of Pcna during the initiation of meiotic recombination when Sycp3 was shown adjacent to the telomeres (Imai et al., 2021). During the leptotene stage, telomeres moved to one pole of the nucleus where Sycp3 aggregated and the accumulated (Saito et al., 2014). The previous and our study suggest that the telomeres serve as a hub where Sycp3, Pcna, MutLγ complex and other factors work coordinately at one pole of the nucleus to initiates meiotic recombination (Blokhina et al., 2019).

## 5. Conclusions

Ddx4, Piwil1, Pcna and Sycp3 have their specific expression pattern in the spermatogenic cells at different differentiation stages. SPG-A can be identified by the highest expression of Ddx4 and Piwil1 among different spermatogenic cells. SPG-B is identified by the second highest expression of Ddx4 and the highest expression of Pcna among different spermatogenic cells. SPD can be distinguished from SPZ by Piwil1 expression. The different subtypes of SPC-I can be identified through co-staining of Sycp3 and Pcna. Leptotene SPC-I expresses both Sycp3 and Pcna polarly at the same side in the nucleus. The antibodies against zebrafish antigen may be used in other fish species. The antibodies along with the integrated method for high-quality immunofluorescence provide a reliable pipeline for spermatogenesis evaluation in fish.

## Abbreviations

SPG: spermatogonia
SPG-A: type-A spermatogonia
SPG-B: type-B spermatogonia
SPC-I: primary spermatocyte
SPC-II: secondary spermatocyte
SPD: spermatid
SPZ: spermatozoa
CZRC: the China Zebrafish Resource Center
RT: room temperature

## Acknowledgement

We thank Kuoyu Li from CZRC for fish care. We thank Fang Zhou from the Analysis and Testing Center of Institute of Hydrobiology for the technical support of confocal microscopy.

This work was supported by the National Key R&D Program of China (2018YFD0901205), National Natural Science Foundation of China (31872550), Strategic Priority Research Program of the Chinese Academy of Sciences (XDA24010108) and the State Key Laboratory of Freshwater Ecology and Biotechnology (grant No. 2019FBZ05).

## Authors Contributions

**Ding Ye**: Project administration, Confocal microscopy, Writing-original draft, Funding acquisition; **Tao Liu**: Investigation, Methodology; **Yongming Li**: Resources; **Yonghua Sun**: Supervision, Writing-review and editing, Funding acquisition

